# Automated assembly scaffolding elevates a new tomato system for high-throughput genome editing

**DOI:** 10.1101/2021.11.18.469135

**Authors:** Michael Alonge, Ludivine Lebeigle, Melanie Kirsche, Sergey Aganezov, Xingang Wang, Zachary B. Lippman, Michael C. Schatz, Sebastian Soyk

## Abstract

Advancing crop genomics requires efficient genetic systems enabled by high-quality personalized genome assemblies. Here, we introduce RagTag, a toolset for automating assembly scaffolding and patching, and we establish chromosome-scale reference genomes for the widely used tomato genotype M82 along with Sweet-100, a rapid-cycling genotype that we developed to accelerate functional genomics and genome editing. This work outlines strategies to rapidly expand genetic systems and genomic resources in other plant species.

## Main

Recent technological advances in genome sequencing and editing enable interrogation and manipulation of crop genomes with unprecedented accuracy. Pan-genomes can capture diverse alleles within crop species but studying their phenotypic consequences is limited by efficient functional genetic systems in relevant and diverse genotypes. Tomato is a model crop system to study the genetics of domestication and quantitative traits. Sequencing hundreds of tomato genomes has uncovered vast genomic diversity [1,2]; however, chromosome-scale genomes are only available for a few accessions [3–5], and there is a historical discrepancy between the reference genome (Heinz 1706) and genotypes that are commonly used for genetic and molecular experimentation (e.g. cultivars M82, Moneymaker, Ailsa Craig, etc.). The large-fruited cultivar M82 has been adopted as a primary reference for genetic, metabolic, and developmental analyses [6,7]; however, a high-quality genome assembly has been lacking, leading to reference bias and false signals in genomics analyses. Furthermore, phenotyping cultivars with larger fruits is labor-intensive and requires extensive growth facilities to accommodate large plants with long generation times. The ultra-dwarfed and small-fruited variety Micro-tom overcomes some of these limitations [8], but a highly mutagenized background, severe hormonal and developmental abnormalities, and low fruit quality undermine its value for studying many phenotypes of translational agronomic importance, such as shoot, inflorescence, and fruit development (**Fig. 1a and Supplementary Fig. 1a-f**).

**Fig. 1.**
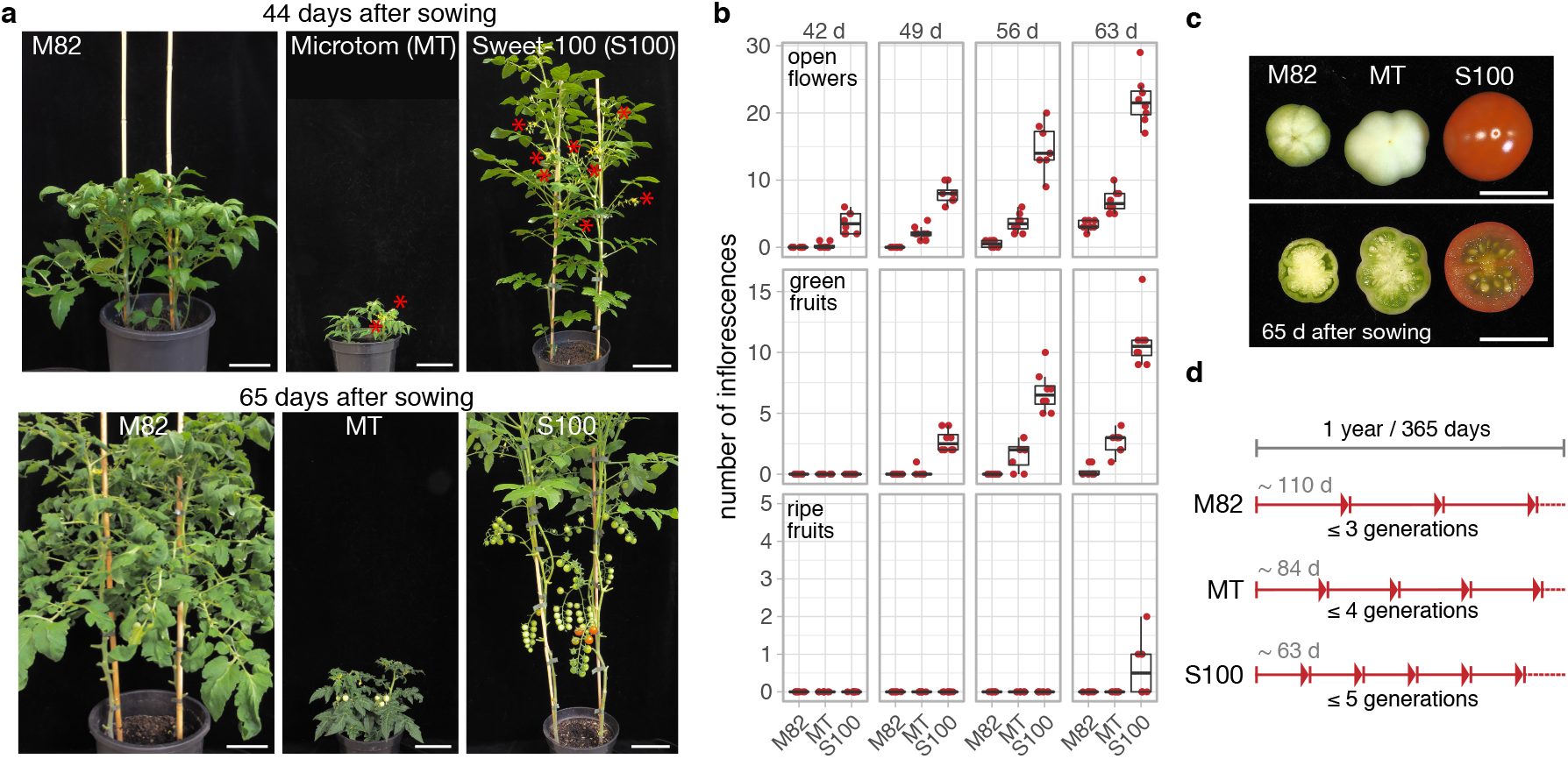
The tomato cultivar Sweet-100 shows characteristics of an efficient experimental system for crop functional genomics. **a**, Images of mature M82, Micro-tom (MT), and Sweet-100 (S100) plants 44 days (top) and 65 days (bottom) after sowing. Red asterisks indicate open flowers. **b**, Number of inflorescences with open flowers (top), green fruits (middle), and ripe fruits (bottom) at 6 to 9 weeks after sowing. Data points represent individual plants. **c**, Images of the first developing fruit on M82, MT, and S100 at 65 days after sowing. **d**, Schematic representation of the generation times of the M82, MT, and S100 genotypes. Scale bars indicate 10 cm (a) and 1 cm (c).

To address these limitations and illustrate how new genomic and genome-editing systems can be rapidly developed, we established the small-fruited tomato cultivar Sweet-100 (S100) as a new system for genome editing and functional genomics. Previously, we used clustered regularly interspaced short palindromic repeat (CRISPR)-Cas9 genome editing to engineer mutations in the paralogous flowering repressor genes *SELF PRUNING* (*SP*) and *SELF PRUNING 5G (SP5G*) in S100 to induce fast flowering and compact growth (**Fig. 1a,b**) [9]. Null mutations in these genes cause rapid-cycling and compact growth without severe developmental abnormalities (**Fig. 1a and Supplementary Fig. 1a-g**). The first ripe fruits mature in only ~65 days after sowing, allowing up to five generations per year compared to at most three generations for most large-fruited genotypes, which also require more space and resources (**Fig. 1a-d**). The short generation time and compact growth habit of S100 allows both greenhouse and field growth at double the normal density with reduced input. Notably, S100 yields ripe fruits ~3 weeks earlier than Micro-Tom and produces more seeds per fruit (**Figure 1b and Supplementary Fig. 1d,f**). Together, these characteristics make S100 a highly efficient system for genetics and a valuable complement to the widely used M82 cultivar. However, a corresponding high-quality reference genome assembly is required to actualize the utility of this new S100 model system.

Genome assemblies are typically built from PacBio High Fidelity (HiFi) and/or Oxford Nanopore long-reads (ONT) [10]. HiFi reads average 15 kbp in length, are highly accurate (~0.1% error), and can produce contiguous draft genome assemblies [11]. However, HiFi-based assemblies often fragment at large and homogenous repeats as well as known sequence-specific coverage dropouts [12]. Built from much longer, though noisier reads with a distinct error-profile, ONT-based assemblies can complement HiFi-based assemblies by resolving some larger repeats or compensating for HiFi coverage dropouts [12]. However, even when using complementary long-read technologies, modern draft genome assemblies rarely achieve complete chromosome-scale. Longer and ultimately chromosome-scale sequences are produced by scaffolding, the process of ordering and orienting genome assembly contigs, and placing gaps between adjacent contigs. Scaffolding is usually achieved by comparing a genome assembly to genome maps encoding the relative distances of genomic markers along chromosomes. Linkage, physical (including optical maps and reference assemblies), and spatial proximity maps (from Chromatin Conformation Capture, or Hi-C data) are popular and effective for scaffolding assemblies. However, because genome maps are noisy and scaffolding methods are fallible, automated scaffolding usually results in incomplete or misassembled scaffolds and researchers often rely on laborious manual curation to correct these shortcomings [13,14].

To help overcome these limitations, we developed RagTag, a new method to automate scaffolding and improve modern genome assemblies (**Fig. 2a**). RagTag supersedes our previously published RaGOO scaffolder by implementing improvements to the homology-based correction and scaffolding modules (see **Methods**) [15] and by providing two new scaffolding tools called “patch” and “merge”. RagTag “patch” uses one genome assembly to make scaffolding joins and fill gaps in a second genome assembly (**Fig. 2b**). This is especially useful for genome assembly projects with complementary sequencing technology types, such as HiFi and ONT, as we demonstrate by accurately patching the CHM13 “Telomere-2-Telomere” human reference assembly (**Extended Data 1 and 2**)(see **Methods**) [12]. [12]. RagTag “merge” is an extension of the CAMSA scaffolder that reconciles multiple candidate scaffolds for a given assembly (**Fig. 2c**) [16]. This allows users to scaffold an assembly with potentially several map or map-specific technical parameters and then synergistically combine results into a single scaffolding solution. The input scaffolds are encoded within a “scaffold graph” with contigs as nodes and edges weighted by the confidence of the placement of contigs. RagTag analyzes this graph to resolve ambiguous paths, optionally using Hi-C data to re-weight the edges [17].

**Fig. 2.**
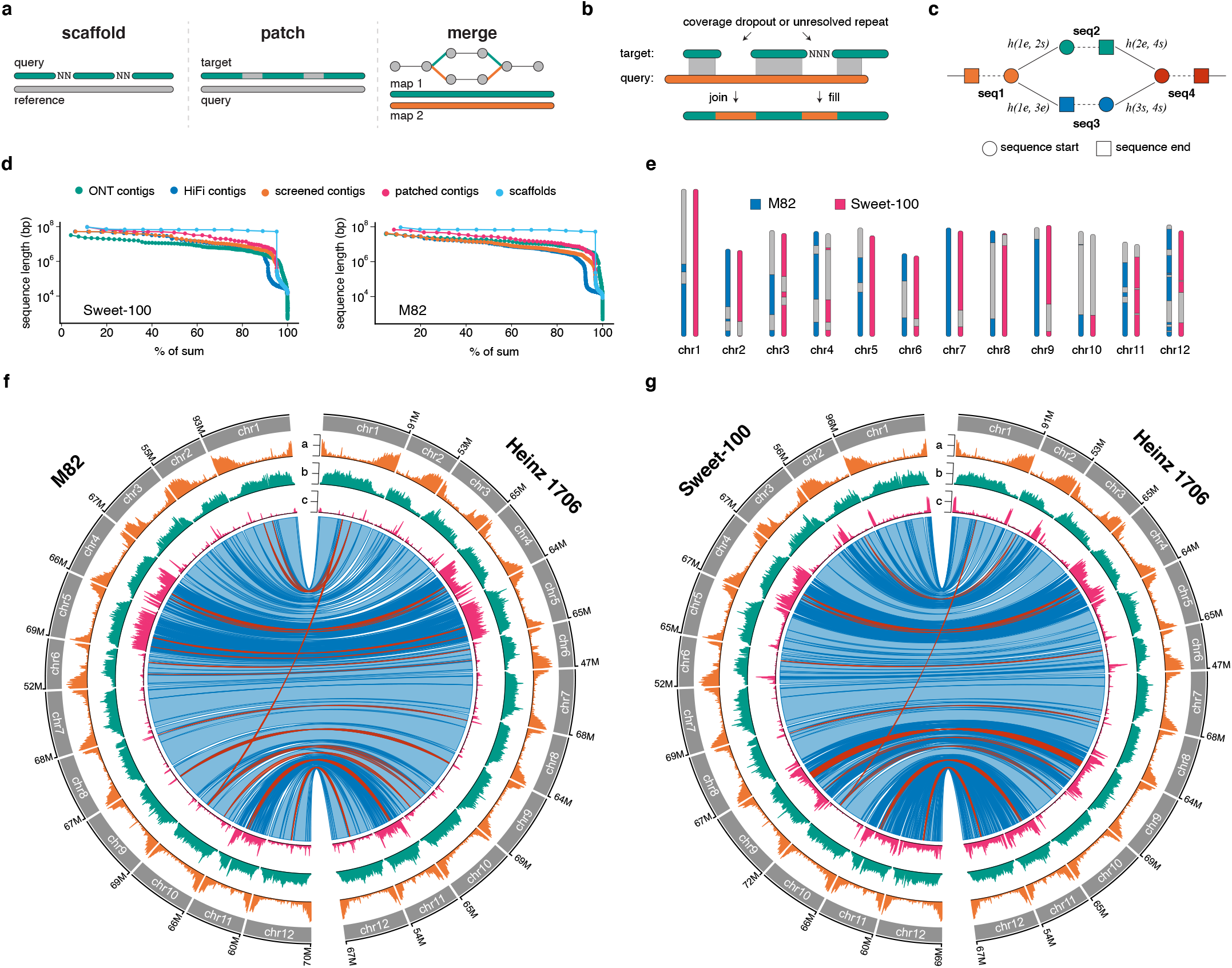
RagTag enables new reference genomes for Sweet-100 and M82. **a**, Overview of RagTag “scaffold”, “patch”, and “merge”. **b**, A more detailed diagram describing RagTag “patch”, highlighting how sequence from the query assembly (orange) can be used to fill gaps in the target assembly (green). **c**, A more detailed diagram describing RagTag “merge” showing how each contig is represented by a pair of nodes for the beginning and end termini of the sequence with edges between contigs indicating the pair of contigs are adjacent in one of the candidate scaffolds. The function *h()* maps contig terminus pairs to Hi-C scores (see **Methods**). **d**, nX plots showing the minimum sequence length (y-axis, log scale) needed to constitute a particular percentage of the assembly (x-axis). **e**, Ideogram showing contig boundaries (alternating color and gray) within the final scaffolds. **f**, Circos plots comparing M82 to Heinz 1706 (SL4.0). Circos quantitative tracks a, b and c are summed in 500 kbp windows and show number of genes (a, lower tick=0, middle tick=47, upper tick=94), LTR retrotransposons (b, 0, 237, 474) and structural variants (c, 0, 24, 48). The inner ribbon track shows wholegenome alignments, with blue indicating forward-strand alignments and red indicating reverse-strand alignments (inversions). Darker colors indicate alignment boundaries. **g**, As for **f** but comparing Sweet-100 to Heinz 1706 and showing number of genes (a, 0, 48, 96), LTR retrotransposons (b, 0, 269, 538) and structural variants (c, 0, 30, 59) and whole-genome alignment ribbons.

We used RagTag to produce high-quality chromosome-scale reference genomes for M82 and S100. Briefly, for each genotype we assembled HiFi and ONT data independently. After screening the HiFi primary contigs for contaminates and organellar sequences, we used RagTag “patch” to patch the HiFi contigs with the ONT contigs, ultimately increasing the N50 from 20.1 to 40.8 Mbp and 12.6 to 27.8 Mbp in S100 and M82, respectively, without introducing any gaps (**Fig. 2d, Supplementary Table 1**). After patching, S100 chromosomes 1 and 5 and M82 chromosome 7 were each represented in a single chromosome-scale contig (**Fig. 2e**). To build 12 chromosome-scale pseudomolecules for each assembly, we incorporated a variety of physical and spatial proximity maps into RagTag “merge” to reconcile alternative scaffolding possibilities into final assembly solutions. Finally, each set of scaffolds were manually validated and corrected with Juicebox Assembly Tools and the assemblies were packaged according to our Pan-Sol specification (https://github.com/pan-sol/pan-sol-spec) [18]. The final S100 and M82 assemblies had a QV score of 56.6 and 53.1, respectively, and re-mapping Hi-C data indicated broad structural accuracy for each assembly (**Supplementary Fig. 2**), demonstrating that RagTag allows fast and accurate generation of chromosome-scale assemblies with little manual intervention.

Using a read mapping approach, we previously reported that S100 and M82 are admixed due to historical breeding, and are thus structurally distinct from the Heinz 1706 reference genome [2]. When comparing all three genomes, we confirmed elevated rates of structural variation across broad chromosomal regions, indicating introgressions from wild relatives (**Fig. 2f, g**). Multiple chromosomes, such as chromosomes 4, 9, 11, and 12 in S100 and chromosomes 4, 5, and 11 in M82, are nearly entirely introgressed from wild relatives, and within introgressions we detected several large inversions. The largest inversion, a ~8.6 Mbp inversion observed on chromosome 9 of S100, was recently discovered in the wild tomato *S. pimpinellifolium* accession LA2093 and was genotyped in 99% of *S. pimpinellifolium* accessions, reinforcing the contribution of *S. pimpinellifolium* and other wild tomato species to the S100 and M82 genomes [4]. Such widespread structural variation between these three tomato accessions highlights the need for personalized genomes to mitigate reference bias and false signals in genomics experiments.

Powerful experimental systems for genetics and functional genomics allow routine genetic manipulation. Using the new S100 genome assembly as a foundation, we adapted our tomato transformation and genome editing protocols to genetically modify S100 (see **Methods**). We obtained transgenic plants in less than four months, comparable to previously published protocols for tomato (**Supplementary Fig. 3a,b**) [8,19]. To test the efficiency of CRISPR-Cas9 genome editing in S100, we utilized our new S100 assembly for accurate guide-RNA (gRNA) design and targeted the tomato homolog of Arabidopsis *APETALA3* (*SlAP3*, Solyc04g081000) on the chromosome 4 introgression with two gRNAs (**Fig. 3a**). In Arabidopsis, *AP3* activity is essential for petal and stamen development [20] and we observed the expected abnormal or missing petals and stamens on all seven *ap3^CR^* first-generation (T0) transgenic plants (**Fig. 3b**). We identified multiple *ap3^CR^* mutant alleles by Sanger sequencing and observed germline transmission in the next generation, demonstrating efficient and robust editing in S100 (**Fig. 3c, d and Supplementary Fig. 3c-e**). We next explored the ability to delete entire gene loci, which is important to mitigate potential confounding genetic compensation responses to mutant allele transcripts [21]. We targeted the floral identity gene *ANANTHA* (*AN*) [22] with two gRNAs 183 bp upstream and 18 bp downstream of the protein coding sequence (**Fig. 3e**). From four T0 transgenics we identified a complete 1568 bp gene deletion allele (*an^CR-1568^*), which was transmitted to the next generation (**Fig. 3f-h, Supplementary Fig. 3f**). Second-generation individuals that carried the *an^CR-1568^* allele developed cauliflower-like inflorescences due to floral meristem overproliferation that is characteristic of *an* mutants. (**Fig. 3h**). Finally, we tested the potential for mutating multigene families and targeted the three floral regulator genes *JOINTLESS2* (*J2*), *ENHANCER OF J2* (*EJ2*), and *LONG INFLORESCENCE* (*LIN*), which belong to the MADS-box gene family [9] (**Fig. 3i**). From eight T0 transgenics, we identified an individual (T0-6) that displayed the *j2^CR^ej2^CR^* double mutant phenotype with highly branched inflorescences, and one individual (T0-17) with the *j2^CR^ej2^CR^lin^CR^* triple mutant null phenotype of cauliflower-like inflorescences (**Fig. 3j**). Sanger sequencing revealed *j2^CR^* and *ej2^CR^* mutant alleles in addition to wild-type *LIN* alleles in the T0-6 individual, while only mutant alleles for all three genes were detected in the T0-17 plant (**Supplementary Fig. 3g**). We then applied a multiplexed amplicon sequencing approach (see **Methods**) and genotyped a population of 184 T1 individuals for edits within protospacer sequences to identify multiple allelic combinations of *j2^CR^*, *ej2^CR^* and *lin^CR^* mutations (**Fig. 3i**). Together, these results illustrate the effectiveness of S100 as an experimental platform for CRISPR-Cas9 genome editing of individual genes and multigene families for functional analyses.

**Fig. 3.**
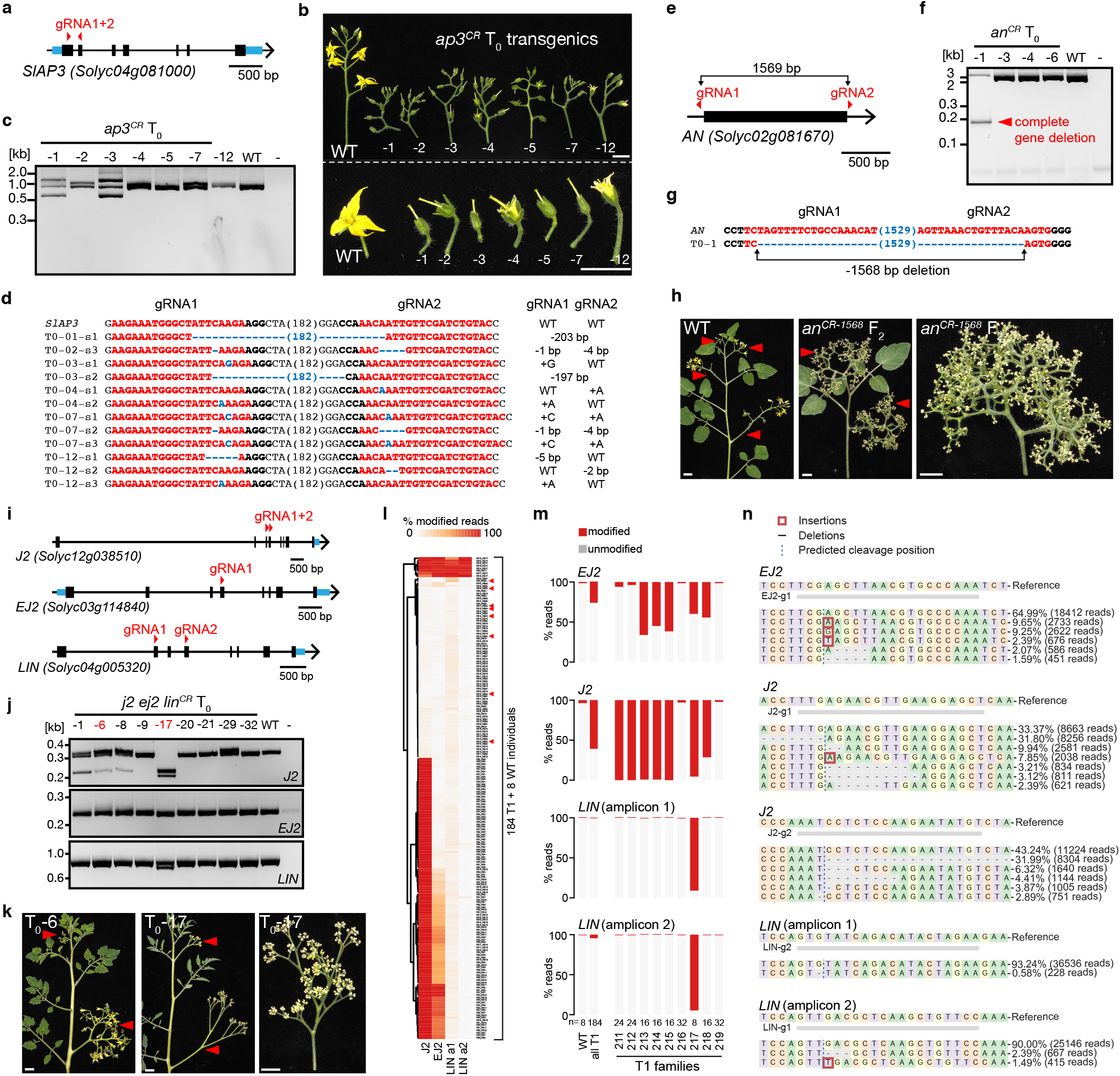
Assessment of genome editing capabilities in Sweet-100. **a**, CRISPR-Cas9 targeting of *SlAP3* using two gRNAs. Black boxes, black lines, and blue boxes represent exonic, intronic, and untranslated regions, respectively. **b**, Images of detached inflorescences (top) and flowers (bottom) from wild-type (WT) and seven independent first-generation (T0) *ap3^CR^* transgenic plants. **c and d**, CRISPR-induced mutations in *SlAP3* identified by agarose gels (c) and Sanger sequencing (d). gRNA and PAM sequences are indicated in red and black bold letters, respectively; deletions are indicated with blue dashes; sequence gap length is given in parenthesis. **e**, Full gene deletion of *AN* by CRISPR-Cas9 using two gRNAs. **f** and **g**, Detection of complete deletion of the *AN* gene by agarose gel electrophoresis (f) and Sanger sequencing (g). **h**, Images of WT and *an^CR^* mutant plants in the non-transgenic second (F2) generation. **i**, CRISPR-Cas9 targeting of the *SEP4* gene family using five gRNAs. **j**, analysis of *j2 ej2 lin^CR^* T0 plants by agarose gel electrophoresis, **k**, images of T0 plants showing *j2 ej2^CR^* double (T0-6) and *j2 ej2 lin^CR^* triple (T0-17) mutant phenotypes. **l**, High-throughput discovery of CRISPR-Cas9 mutations in *J2*, *EJ2*, and *LIN* by multiplexed amplicon sequencing. Heatmap shows the percentage of modified reads in 184 T1 and 8 WT control plants. Red arrowheads indicate WT control individuals. **m**, Quantification of editing efficiency (percentage of modified reads) in WT, all T1, and individual T1 families; *n* equals the number of individual plants. **n**, Sequences and frequency of edited alleles identified from in the T1 generation. Scale bars indicate 1 cm.

In summary, we introduced a new toolset for automated genome assembly scaffolding and we elevated a rapid-cycling tomato variety to an effective experimental system for functional genomics and genome-scale editing experiments. Our study outlines a roadmap to rapidly establish multiple personalized reference systems as cornerstones for functional interrogation of the vast genetic variation within and between species.

## Methods

### Plant material, growth conditions, and phenotyping

Seeds of *S. lycopersicum* cv. M82 (LA3475), Sweet-100 (S100), and Micro-tom (MT) were from our own stocks. Seeds were directly sown and germinated in soil in 96-cell plastic flats. Plants were grown under long-day conditions (16-h light, 8-h dark) in a greenhouse under natural light supplemented with artificial light from high-pressure sodium bulbs (~250 μmol m^−2^ s^−1^) at 25°C and 50-60% relative humidity. Seedlings were transplanted to soil to 3.5 l (S100 and MT) or 10 l (M82) pots 3-4 weeks after sowing. Analyses of fruit ripening, flower number, seed number, fruit weight, fruit sugar content (Brix), and inflorescence branching were conducted on mature plants grown in pots. Sugar content (Brix) of fruit juice was quantified using a digital refractometer (Hanna Instruments HI96811). Fruit ripening was quantified by labeling individual flowers at anthesis and counting the days to breaker fruit stage and red fruit stage.

### RagTag Overview

RagTag supersedes RaGOO as a homology-based genome assembly correction (RagTag “correct”) and scaffolding (RagTag “scaffold”) tool [15]. RagTag implements several general improvements and conveniences for these features but follows the same algorithmic approach as previously reported. RagTag also provides two new tools called “patch” and “merge” for genome assembly improvement. RagTag “patch” uses one genome assembly to “patch” (join contigs and/or fill gaps) sequences in another assembly. RagTag “merge” reconciles two or more distinct scaffolding solutions for the same assembly. Finally, RagTag offers a variety of command-line utilities for calculating assembly statistics, validating AGP files, and manipulating genome assembly file formats. RagTag is open source (distributed under the MIT license) and is available on GitHub: https://github.com/malonge/RagTag

### RagTag whole-genome alignment filtering and merging

Most RagTag tools rely on pairwise (a “query” vs. a “reference/target”) whole-genome alignments. RagTag supports the use of Minimap2, Unimap, or Nucmer for whole-genome alignment, though any alignments in PAF or MUMmer’s delta format can be used [23,24]. RagTag filters and merges whole-genome alignments to extract useful scaffolding information. To remove repetitive alignments, RagTag uses an integrated version of unique anchor filtering introduced by Assemblytics [25]. RagTag can also remove alignments based on mapping quality score, when available. Filtered alignments are then merged to identify macro-synteny blocks. For each query sequence, alignments are sorted by reference position. Consecutive alignments within 100 kbp (configured using the “-d” parameter) of each other and on the same strand are merged together, taking the minimum coordinate as the new start position and the maximum coordinate as the new end position. Consequently, unmerged alignments are either far apart on the same reference sequence, on different reference sequences, or on different strands. Finally, merged alignments contained within other merged alignments (with respect to the query position) are removed.

### RagTag “correct”

Following the approach we developed for RaGOO, RagTag “correct” uses pairwise whole-genome sequence homology to identify and correct putative misassemblies. First, RagTag generates filtered and merged whole-genome alignments between a “query” and a “reference” assembly. The “query” assembly will be corrected and the “reference” assembly will be used to inform correction. Any query sequence with more than one merged alignment is considered for correction. RagTag breaks these query sequences at merged alignment boundaries provided that the boundaries are not within 5 kbp (−b) from either sequence terminus. Users may optionally choose to only break between alignments to the same or different reference sequences (--intra and --inter). If a GFF file is provided to annotate features in the query assembly, the query assembly will never be broken within a defined feature.

When the query and reference assemblies do not represent the same genotypes, unmerged alignments within a contig can indicate genuine structural variation. To help distinguish between structural variation and misassemblies, users can optionally provide Whole Genome Shotgun (WGS) sequencing reads from the same query genotype, such as short accurate reads or long error-corrected reads, to validate putative query breakpoints. RagTag aligns these reads to the query assembly with Minimap2 and computes the read coverage for each position in the query assembly. For each proposed query breakpoint, RagTag will flag exceptionally low (below --min-cov) or high (above --max-cov) coverage within 10 kbp (−v) of the proposed breakpoint. If exceptionally low or high coverage is not observed, the merged alignment boundaries are considered to be caused by true variation, and the query assembly is not broken at this position.

### RagTag “scaffold”

RagTag “scaffold” uses pairwise whole-genome sequence homology to scaffold a genome assembly. First, RagTag generates filtered and merged whole-genome alignments between a “query” and a “reference” assembly. The “query” assembly will be scaffolded and the “reference” assembly will be used to inform scaffolding. The merged alignments are used to compute a clustering, location, and orientation “confidence” score, just as is done in RaGOO, and sequences with confidence scores below certain thresholds are excluded (as set with parameters “-i”, “-a”, and “-s”). For each query sequence, the longest merged alignment is designated as the “primary” alignment. Primary alignments contained within other primary alignments (with respect to the reference coordinates) are removed. Primary alignments are then used to order and orient query sequences. To order query sequences, sequences are assigned to the reference chromosome to which they primarily align. Then, for each reference sequence, primary alignments are sorted by reference coordinate, establishing an order of query sequences. To orient query sequences, the sequence is assigned the same orientation as its primary alignment. Query sequences with no filtered alignments to the reference assembly (“unplaced” sequences) are output without modification or are optionally concatenated together.

By default, 100 bp gaps are placed between adjacent scaffolded query sequences, indicating an “unknown” gap size according to the AGP specification (https://www.ncbi.nlm.nih.gov/assembly/agp/AGP_Specification/). Optionally, RagTag can infer the gap size based on the whole-genome pairwise alignments. Let *seq1* (upstream) and *seq2* (downstream) be adjacent query sequences, and let *aln1* and *aln2* be their respective primary alignments. Let *rs*, *re*, *qs*, and *qe* denote the alignment reference start position, reference end position, query start position, and query end position, respectively. The following function computes the inferred gap length between *seq1* and *seq2*:

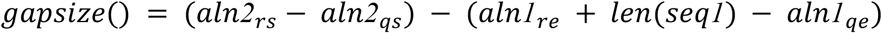

Where *len*(*seq*) is the length of *seq1*. All inferred gap sizes must be at least 1 bp, and if the inferred gap size is too small (-g or less than 1) or too large (-m), it is replaced with an “unknown” gap size of 100 bp.

### RagTag “patch”

The new RagTag “patch” tool uses pairwise whole-genome sequence homology to make joins between contigs, without introducing gaps, and fill gaps in a “target” genome assembly using sequences from a “query” genome assembly. First, RagTag breaks all target sequences at gaps and generates filtered and merged whole-genome alignments between the query and target assemblies. Merged alignments that are not close (-i) to a target sequence terminus or are shorter than 50000 bp (-s) are removed. If an alignment is not close to both query sequence termini yet it is not close to either target sequence terminus, meaning the target sequence should be contained within the query sequence, yet large portions of the target sequence do not align to the query sequence, the alignment is discarded.

To ultimately patch the target assembly, RagTag employs a directed version of a “scaffold graph” [16,26]. Nodes in the graph are target sequence termini (two per target sequence), and edges connect termini of distinct target sequences observed to be adjacent in the input candidate scaffolds. The graph is initialized with the known target sequence adjacencies originally separated by gaps in the target assembly. Next, merged and filtered alignments are processed to identify new target sequence adjacencies. For each query sequence that aligns to more than one target sequence, alignments are sorted by query position. For each pair of adjacent target sequences, an edge is created in the scaffold graph. The edge stores metadata such as query sequence coordinates in order to continuously join the adjacent target sequences. If an edge already exists due to an existing gap, the gap metadata is replaced with the query sequence metadata so that the gap can be replaced with sequence. If an adjacency is supported by more than one alignment, the corresponding edge is discarded. To find a solution to this graph and output a patched assembly, a maximal weight matching is computed with networkx and if there are any cycles, they are broken [27]. RagTag then iterates through each connected component and iteratively builds a sequence from adjacent target sequences. When target sequences are not overlapping, they are connected with sequence from the supporting query sequence. Unpatched target sequences are output without modification.

### RagTag “merge”

RagTag “merge” is a new implementation and extension of CAMSA, a tool to reconcile two or more distinct scaffolding solutions for a genome assembly [16]. Input scaffolding solutions must be in valid AGP format, and they must order and orient the same set of genome assembly AGP “components”. RagTag iteratively builds a scaffold graph to store adjacency evidence provided by each AGP file. First, each AGP file is assigned a weight (1 by default). Then, for each AGP file and for each pair of adjacent components, an edge is added to the scaffold graph, and the edge weight is incremented by the weight of the AGP file, just as is done in CAMSA. After the scaffold graph is created, users can optionally replace native edge weights with Hi-C weights. To do this, Hi-C alignments are used to compute scaffold graph weights according to the SALSA2 algorithm, which uses the same underlying scaffold graph data structure. To find a solution to this graph and to output a merged AGP file, a maximal weight matching is computed with networkx and if there are any cycles, they are broken. RagTag then iterates through each connected component and iteratively builds AGP objects. Unmerged components are output without modification.

### Patching a human Telomere-2-Telomere genome assembly

The CHM13v1.1 assembly is the first-ever published complete sequence of a human genome [12]. Though the original draft assembly was built exclusively from HiFi reads, it was manually inspected and patched at 25 loci, mostly at HiFi coverage dropouts, with sequence from the previously published, ONT-based CHM13v0.7 assembly. Using these 25 manual patches as a benchmark (**Supplementary Data 1**), we evaluated the ability of RagTag to automatically patch the CHM13 draft assembly with the CHM13v0.7 assembly. RagTag made all 25 patches (**Supplementary Data 2**), 19 of which were identical to the manual patches. The remaining six patches had slightly shifted patch coordinates, with a median Euclidean distance of 66.4 bp using the start and end genomic coordinates for the two joined sequences and the sequence used for patching. The slight differences in coordinates are due to locally repetitive sequences that cause aligner-specific coordinates to be selected when transitioning from the query and target sequences. RagTag made one false join connecting chr18 and chr10, though this was caused by a misassembly in CHM13v.07 caused by a long, high-identity repeat shared between these chromosomes. Patching was performed with RagTag v2.1.0 (--aligner minimap2).

### Extraction of high-molecular weight DNA and sequencing

Extraction of high-molecular weight genomic DNA, construction of Oxford Nanopore Technology libraries and sequencing were described previously [2]. Libraries for PacBio HiFi sequencing were constructed and sequenced at the Genome Technology Center at UNIL and Genome Center at CSHL. High-molecular-weight DNA was sheared with a Megaruptor (Diagenode) to obtain 15-20 kb fragments. After shearing, the DNA size distribution was evaluated with a Fragment Analyzer (Agilent) and 5-10 μg of the DNA was used to prepare a SMRTbell library with the PacBio SMRTbell Express Template Prep Kit 2.0 (Pacific Biosciences) according to the manufacturer’s instructions. The library was size-selected on a BluePippin system (Sage Science) for molecules larger than 12.5 kb and sequenced on one SMRT cell 8M with v2.0/v2.0 chemistry on a PacBio Sequel II instrument (Pacific Biosciences) at 30 hours movie length. Hi-C experiments were conducted at Arima Genomics (San Diego, CA) from 2 g of flash-frozen leaf tissue.

### BLAST databases for screening contigs

We built each BLAST database with makeblastdb (v2.5.0+, −dbtype nucl) [28]. We used all RefSeq bacterial genomes (downloaded on February 11th, 2021) for the bacterial genomes database. We used a collection of Solanum chloroplast sequences for the chloroplast database, and their GenBank accession IDs are as follows:

> MN218076.1, MN218077.1, MN218078.1, MN218079.1, MN218091.1, MN218088.1, MN218089.1, NC_039611.1, NC_035724.1, KX792501.2, NC_041604.1, MH283721.1, NC_039605.1, NC_039600.1, NC_007898.3, MN218081.1, NC_039606.1, NC_030207.1, MT120858.1, MN635796.1, MN218090.1, MT120855.1, MT120856.1, NC_050206.1, MN218087.1, NC_008096.2

We used a collection of Solanum mitochondrial sequences for the mitochondria database, and their GenBank accession IDs are as follows:

> MT122954.1, MT122955.1, MT122966.1, MT122969.1, MT122973.1, MT122974.1, MT122977.1, MT122988.1, NC_050335.1, MT122980.1, MT122981.1, MT122982.1, MT122983.1, MF989960.1, MF989961.1, NC_035963.1, MT122970.1, MT122971.1, NC_050334.1, MW122958.1, MW122959.1, MW122960.1, MT122964.1, MT122965.1, MW122949.1, MW122950.1, MW122951.1, MW122952.1, MW122953.1, MW122954.1, MW122961.1, MW122962.1, MW122963.1, MT122978.1, MT122979.1, MF989953.1, MF989957.1, MN114537.1, MN114538.1, MN114539.1, MT122958.1, MT122959.1

We used a collection of Solanum rDNA sequences for the rDNA database, and their GenBank accession IDs are as follows:

> X55697.1, AY366528.1, AY366529.1, KF156909.1, KF156910.1, KF156911.1, KF156912.1, KF156913.1, KF156914.1, KF156915.1, KF156916.1, KF156917.1, KF156918.1, KF156919.1, KF156920.1, KF156921.1, KF156922.1, KF603895.1, KF603896.1, X65489.1, X82780.1, AF464863.1, AF464865.1, AY366530.1, AY366531.1, AY875827.1

### Sweet-100 genome assembly

The following describes the methods used to produce SollycSweet-100_v2.0 assembly. We independently assembled all HiFi reads (33,815,891,985 bp) with Hifiasm (v0.13-r308, −l0) and we assembled ONT reads at least 30 kbp long (a total of 28,595,007,408 bp) with Flye (v2.8.2-b1689, --genome-size 1g) [29,30]. The Hifiasm primary contigs were screened to remove contaminant or organellar contigs using the databases described above. Next we used WindowMasker to mask repeats in the primary contigs (v1.0.0, −mk_counts -sformat obinary - genome_size 882654037) [31]. We then aligned each contig to the bacterial, chloroplast, mitochondria, and rDNA BLAST databases with blastn (v2.5.0+, -task megablast). We only included the WindowMasker file for alignments to the bacterial database (-window_masker_db). For each contig, we counted the percentage of base pairs covered by alignments to each database. If more than 10% of a contig aligned to the rDNA database, we deemed it to be a putative rDNA contig. We then removed any contigs not identified as rDNA contigs that met any of the following criteria: 1) More than 10% of the contig was covered by alignments to the bacterial database; 2) More than 20% of the contig was covered by alignments to the mitochondria database and the contig was less than 1 Mbp long; or 3) More than 20% of the contig was covered by alignments to the chloroplast database and the contig was less than 0.5 Mbp long. In total, we removed 1,015 contigs (35,481,360 bp) with an average length of 34,957.005 bp, most of which contained chloroplast sequence.

Even though Sweet-100 is an inbred line, to ensure that the assembly did not contain haplotypic duplication, we aligned all HiFi reads to the screened Hifiasm contigs with Winnowmap2 (v2.0, k=15, --MD -ax map-pb) [23]. We then used purge_dups to compute and visualize the contig coverage distribution, and we determined that haplotypic duplication was not evident in the screened contigs [32].

We used RagTag “patch” to patch the screened Hifiasm contigs with sequences from the ONT flye contigs, and we manually excluded three incorrect patches caused by a missassembly in the Flye contigs. We then scaffolded the patched contigs using three separate approaches producing three separate AGP files. For the first two approaches, we used RagTag for homology-based scaffolding, once using the SL4.0 reference genome and once using the LA2093 v1.5 reference genome (v2.0.1, --aligner=nucmer --nucmer-params=“--maxmatch -l 100 -c 500”) [3,4]. In both cases, only contigs at least 100 kbp long were considered for scaffolding, and the reference chromosome 0 sequences were not used for scaffolding. For the third scaffolding approach, we used Juicebox Assembly Tools to manually scaffold contigs with Hi-C data (using “arima” as the restriction enzyme), and we used a custom script to convert the “.assembly” file to an AGP file. We also separately generated Hi-C alignments by aligning the Hi-C reads to the screened contigs with bwa mem (v0.7.17-r1198-dirty) and processing the alignments with the Arima mapping pipeline (https://github.com/ArimaGenomics/mapping_pipeline) which employs Picard Tools (https://broadinstitute.github.io/picard/) [33]. We merged the three AGP files with RagTag “merge” (v2.0.1, -r ‘GATC,GA[ATCG]TC,CT[ATCG]AG,TTAA’), using Hi-C alignments to weight the Scaffold Graph (-b). Finally, using the merged scaffolds as a template, we made four manual scaffolding corrections in Juicebox Assembly tools. The final assembly contained 12 scaffolds corresponding to 12 chromosomes totaling 805,184,690 bp of sequence and 918 unplaced nuclear sequences totaling 40,749,555 bp.

VecScreen did not identify any “strong” or “moderate” hits to the adaptor contamination database (ftp://ftp.ncbi.nlm.nih.gov/pub/kitts/adaptors_for_screening_euks.fa)

(https://www.ncbi.nlm.nih.gov/tools/vecscreen/). We packaged the assembly according to the pan-sol v0 specification (https://github.com/pan-sol/pan-sol-spec), and chromosomes were renamed and oriented to match the SL4.0 reference genome. The tomato chloroplast (GenBank accession NC_007898.3) and mitochondria (GenBank accession NC_035963.1) reference genomes were added to the final assembly.

To identify potential misassemblies and heterozygous Structural Variants (SVs), we aligned all HiFi reads (v2.0, k=15, --MD -ax map-pb) and ONT reads longer than 30 kbp (v2.0, k=15, --MD -ax map-ont) to the final assembly with Winnowmap2 and we called structural variants with Sniffles (v1.0.12, −d 50 −n −1 −s 5) [34]. We removed any SVs with less than 30% of reads supporting the ALT allele and we merged the filtered SV calls (317 in total) with Jasmine (v1.0.10, max_dist=500 spec_reads=5 --output_genotypes) [35,36].

### Sweet-100 gene and repeat annotation

We used Liftoff to annotate the Sweet 100 v2.0 assembly using ITAG4.0 gene models and tomato pan-genome genes as evidence (v1.5.1, -copies) [1,3,37]. Chloroplast and mitochondria annotations were replaced with their original GenBank annotation. Transcript, coding sequence, and protein sequences were extracted using gffread (v0.12.3, -y -w -x) [38]. We annotated transposable elements with EDTA (v1.9.6, --cds --overwrite 1 --sensitive 1 --anno 1 --evaluate 1) [39].

### M82 genome assembly

The M82 genome was assembled following the approach used for the Sweet-100 assembly, with the following distinctions. First, Hifiasm v0.15-r327 was used for assembling HiFi reads. Also, the M82 ONT assembly was polished before patching. M82 Illumina short-reads [15] were aligned to the draft Flye ONT assembly with BWA-MEM (v0.7.17-r1198-dirty) and alignments were sorted and compressed with samtools (v1.10) [33,40]. Small variants were called with freebayes (v1.3.2-dirty, --skip-coverage 480) and polishing edits were incorporated into the assembly with bcftools “consensus” (v1.10.2, -i’QUAL>1 && (GT=“AA” || GT=“Aa”)’ -Hla) [41]. In total, two iterative rounds of polishing were used. RagTag “merge” was also used for scaffolding, though the input scaffolding solutions used different methods than the Sweet-100 assembly. First, homology-based scaffolds were generated with RagTag “scaffold”, using the SL4.0 reference genome (v2.0.1, --aligner=nucmer --nucmer-params=“--maxmatch -l 100 -c 500”). Contigs smaller than 300 kbp were not scaffolded (-j), and the reference chromosome 0 was not used to inform scaffolding (-e). Next, SALSA2 was used to derive Hi-C-based scaffolds. Hi-C reads were aligned to the assembly with the pipeline described for Sweet-100. We then produced scaffolds with SALSA2 (-c 300000 -p yes -e GATC -m no) and manually corrected false scaffolding joins in Juicebox Assembly Tools. We reconciled the homology-based and Hi-C-based scaffolds with RagTag “merge” using Hi-C alignments to re-weight the scaffold graph (-b). Finally, we made four manual corrections in Juicebox Assembly Tools. Cooler and HiGlass were used to visualize Hi-C heatmaps [42,43]. Mercury was used to calculate QV and k-mer completeness metrics [44].

### CRISPR-Cas9 mutagenesis, plant transformation, and identification of mutant alleles

CRISPR-Cas9 mutagenesis was performed as described previously [45]. Briefly, guide RNAs (gRNAs) were designed based on the Sweet 100 v2.0 assembly and the CRISPRdirect tool (https://crispr.dbcls.jp/). Binary vectors for plant transformation were assembled using the Golden Gate cloning system as previously described [46]. Final vectors were transformed into the tomato cultivar S100 by *Agrobacterium tumefaciens-mediated* transformation according to Gupta and Van Eck (2016) with minor modifications [19]. Briefly, seeds were sterilized for 15 min in 1.3% bleach followed by 10 min in 70% ethanol and rinsed four times with sterile water before sowing on MS media (4.4 g/l MS salts, 1.5 % sucrose, 0.8 % agar, pH 5.9) in Magenta boxes. Cotyledons were excised 7-8 days after sowing and incubated on 2Z-media [19] at 25°C in the dark for 24 hrs before transformation. *A. tumefaciens* were grown in LB media and washed in MS-0.2% media (4.4 g/l MS salts, 2% sucrose, 100 mg/l myo-inositol, 0.4 mg/l thiamine, 2 mg/l acetosyringone, pH5.8). Explants were co-cultivated with *A. tumefaciens* on 2Z-media supplemented with 100 μg/l IAA for 48 hrs at 25°C in the dark and transferred to 2Z selection media (supplemented with 150 mg/l kanamycin). Explants were transferred every two weeks to fresh 2Z selection media until shoot regeneration. Shoots were excised and transferred to selective rooting media [19] (supplemented with 150 mg/l kanamycin) in Magenta boxes. Well-rooted shoots were transplanted to soil and acclimated in a Percival growth chamber (~100 μmol m^−2^ s^−1^, 25°C, 50% humidity) before transfer to the greenhouse.

Genomic DNA was extracted from T0 plants using a quick genomic DNA extraction protocol. Briefly, small pieces of leaf tissue were flash frozen in liquid nitrogen and ground in a bead mill (Qiagen). Tissue powder was incubated in extraction buffer (100 mM Tris-HCl pH9.5, 250 mM KCl, 10 mM EDTA) for 10 min at 95°C followed by 5 min on ice. Extracts were combined with one volume of 3% BSA, vigorously vortexed, and spun at 13.000 rpm for 1 min. One microliter supernatant was used as template for PCR using primers flanking the gRNA target sites. PCR products were separated on agarose gels and purified for Sanger Sequencing (Microsynth) using ExoSAP-IT reagent (NEB). Chimeric PCR products were subcloned before sequencing using StrataClone PCR cloning kits (Agilent).

High-throughput genotyping of T1 individuals was conducted by barcoded amplicon sequencing according to Liu et al., 2021 with minor modifications [47]. Briefly, gene-specific amplicons were diluted ten-fold before barcoding and pools of barcoded amplicons were gel-purified (NEB Monarch DNA gel extraction) before Illumina library preparation and sequencing (Amplicon-EZ service at Genewiz). Editing efficiencies were quantified from a total of 194,590 aligned reads using the CRISPResso2 software (--min_frequency_alleles_around_cut_to_plot 0.1 --quantification_window_size 50) [48]. All oligos used in this study are listed in **Supplementary Tables 2-4**.

## Data availability

Genome assemblies and annotations are available at https://github.com/pan-sol/pan-sol-data. Raw sequence data is available on SRA under the BioProject PRJNA779684. Source Data files for all main and supplementary figures will be available in the online version of the paper. Seeds are available on request from S. Soyk. All additional data sets are available from the corresponding author on request.

## Code availability

The RagTag software is freely available on GitHub under the MIT license: https://github.com/malonge/RagTag

## Acknowledgments

We thank all members of the Lippman, Schatz, and Soyk labs for helpful discussions. We thank B. Tissot-Dit-Sanfin, A. Robadey, and V. Vashanthakumar from UNIL for assistance with plant care. We thank J. Marquis, J. Weber, and M. Dupasquier from the UNIL Genome Technology Facility (GTF) for sequencing support and René Dreos for bioinformatic support. We thank Sara Goodwin, Melissa Kramer, and the Cold Spring Harbor Laboratory Woodbury Genome Center for sequencing support. We thank Lei Liu and Penelope Lindsay for helpful discussions about amplicon sequencing. We thank Steven Salzberg, Aleksey Zimin, Luca Venturini, and Alexander Leonard for their help with software features and improvements. This work was performed with assistance from the US National Institutes of Health Grant S10OD028632-01. M.A. and M.C.S. were supported by the National Science Foundation (NSF) DBI-1350041, and NSF PGRP grant IOS-1732253 to Z.B.L. and M.C.S. This work was supported by funding from the Howard Hughes Medical Institutes to Z.B.L., and the European Research Council (ERC) under the European Union’s Horizon 2020 research and innovation programme (ERC Starting Grant “EPICROP” Grant No. 802008) and a Swiss National Science Foundation (SNSF) Eccellenza Professorial Fellowship (Grant No. PCEFP3_181238) to S.S.

## Author contributions

M.A., Z.B.L, M.C.S., and S.S. conceived the project and designed and planned experiments. M.A., L. L., Z.B.L, M.C.S., and S.S. performed experiments and collected data. M.A., L.L., X.W., Z.B.L, M.C.S., and S.S. analyzed data. M.A., M.K., and S.A. designed the RagTag software. M.A. developed the RagTag software. M.A., and S.S. wrote the manuscript with editing from Z.B.L. and M.C.S. All authors read, edited, and approved the manuscript.

---

**Supplementary Fig. 1.**
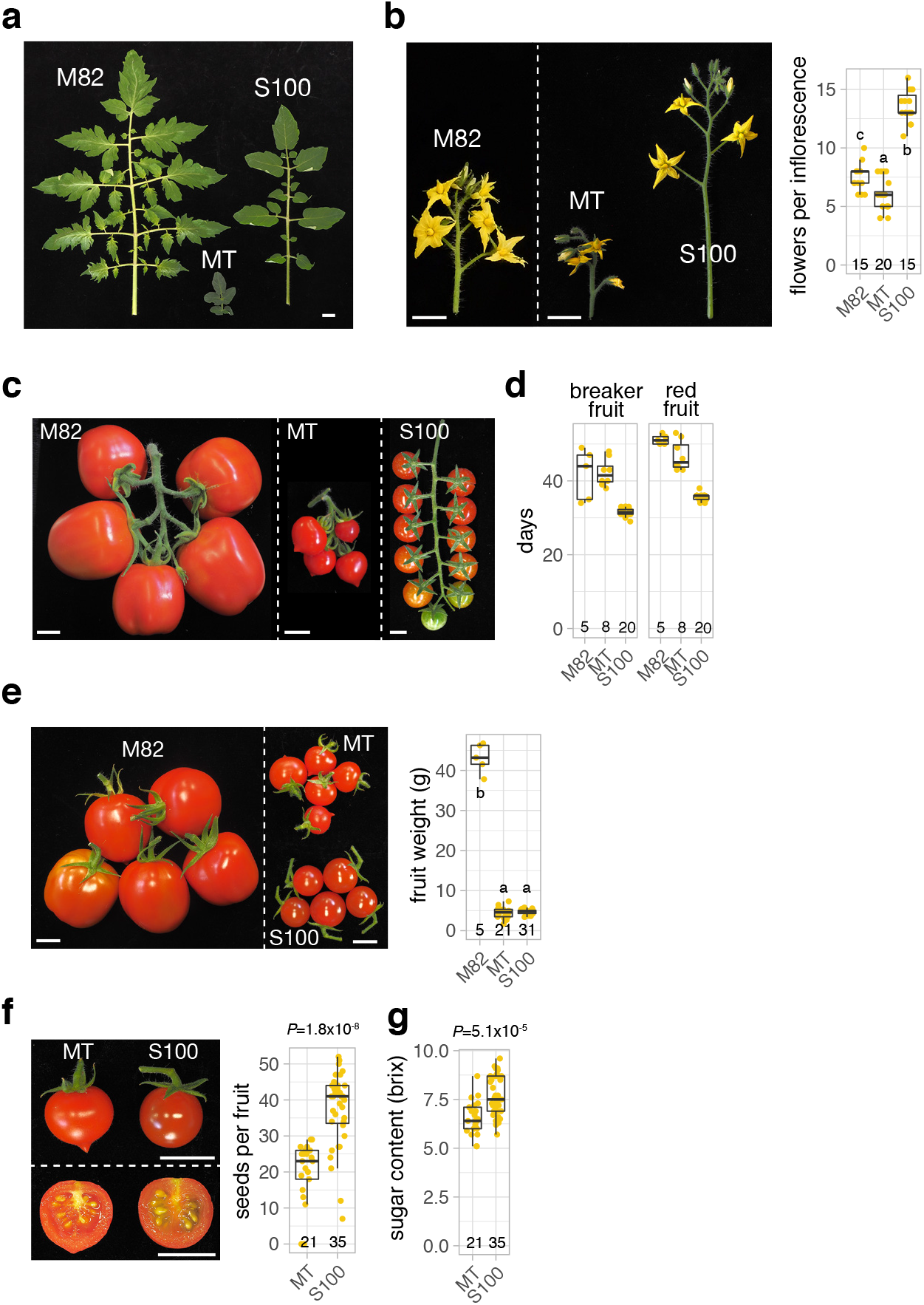
Phenotypic characteristics of the tomato cultivars M82 and Sweet-100 (S100), and the experimental variety Micro-tom (MT). **a**, Images of mature leaves from M82, MT, and S100. **b**, Images of detached inflorescences from M82, MT, and S100 plants and quantification of flower number per inflorescence. n equals the number of inflorescences. **c,** Images of detached fruit clusters from M82, S100 and MT. **d**, Quantification of days from anthesis to breaker fruit (left) and red fruit (right panel) in M82, MT, and S100. **e,** Images of fruits from M82, MT, and S100 and quantification of fruit weight. **f**, Images of fruits and quantification of seed number per fruit in MT and S100. n equals the number of fruits. **g**, Quantification of fruit sugar content in MT and S100. n in d-g equals the number of fruits. Scale bars indicate 1 cm. Letters in c and e represent post-hoc Tukey’s HSD tests. *P* values in f and g represent two-tailed, two-sample *t*-tests.

**Supplementary Fig. 2.**
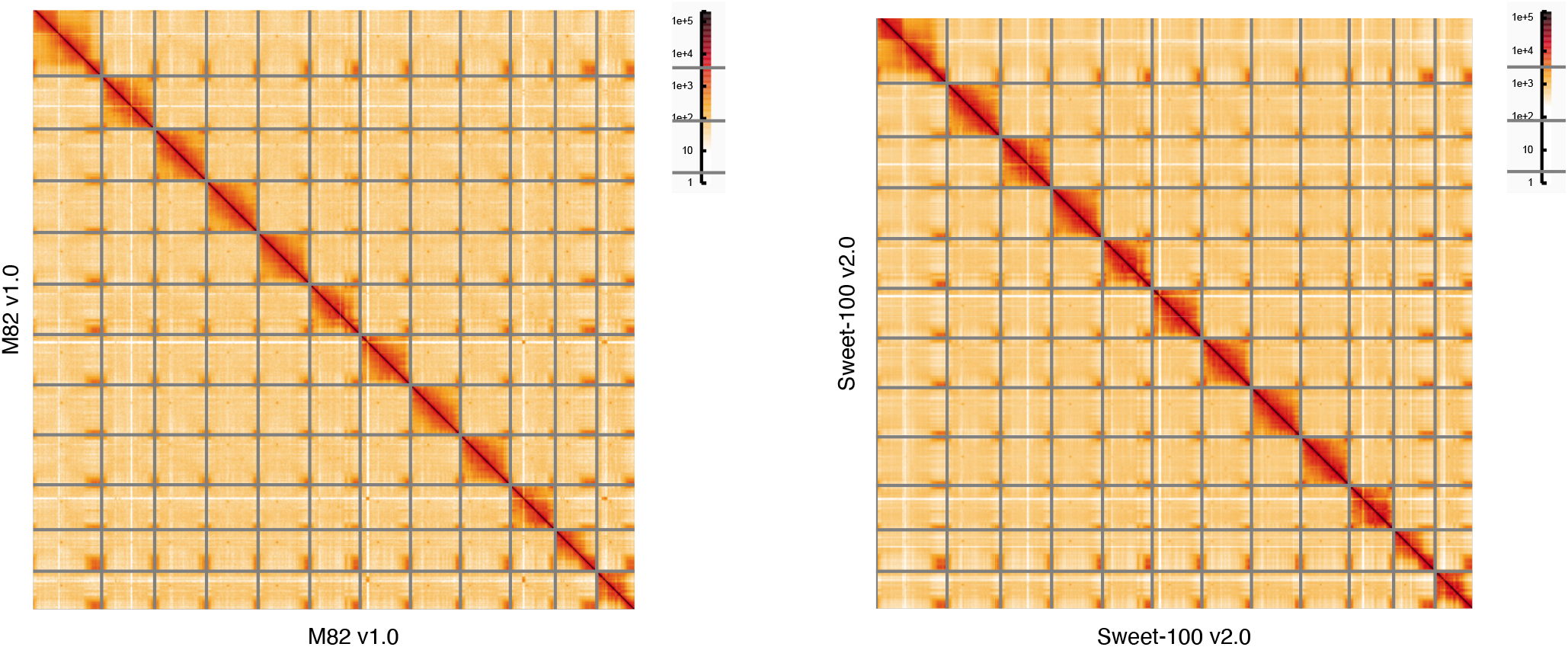
Hi-C heatmaps confirm reference assembly structural accuracy. Hi-C heatmaps for the M82 v1.0 and the Sweet-100 v2.0 reference assemblies. The 12 chromosomes are sorted from largest (top left) to smallest (bottom right).

**Supplementary Fig. 3.**
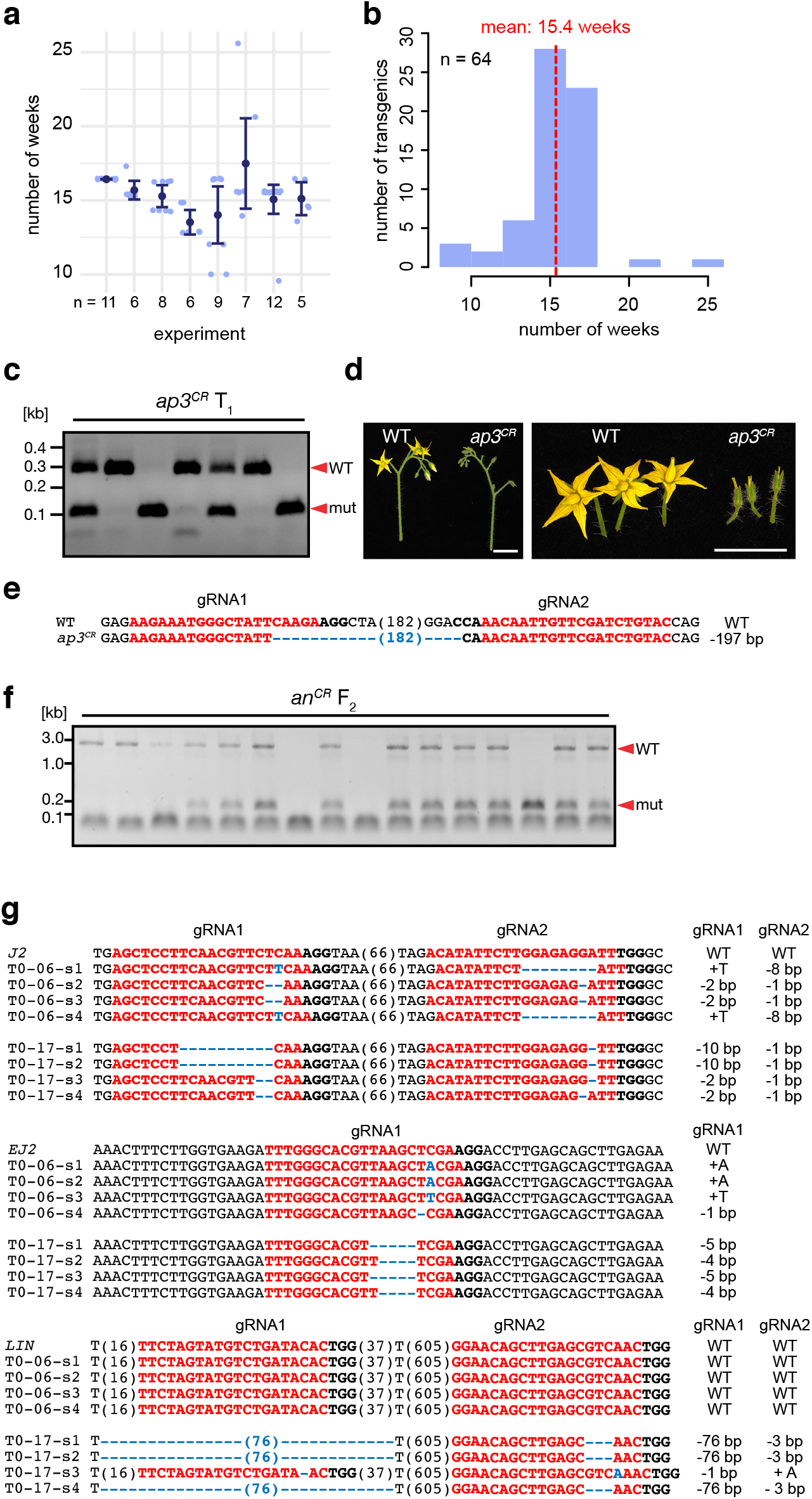
Plant transformation and CRISPR-Cas9 genome editing in Sweet-100. **a and b**, Number of weeks for obtaining transgenic plants, determined in weeks from sowing to transfer to soil. Data is shown for eight independent transformation experiments. n equals number of transgenic plants. **c**, PCR-genotyping for deletions in *SlAP3* in the non-transgenic second (F2) generation. **d**, Images of detached inflorescences (left) and flowers (right) from wild-type (WT) and *ap3^CR^* F2 plants. **e**, Sequence of CRISPR-induced *ap3^CR^* allele transmitted to *ap3^CR^* F2 plants. gRNA and PAM sequences are indicated in red and black bold letters, respectively; deletions are indicated with blue dashes; deletions; sequence gap length is given in parenthesis. **f**, PCR-genotyping for deletions in *AN* in the non-transgenic second (F2) generation. **g**, CRISPR-induced mutations in *J2*, *EJ2*, and *LIN* identified by Sanger sequencing of T0 plants. Scale bars indicate 1cm.

